# The relationship between task-related aperiodic EEG activity, neural inefficiency and verbal working memory in younger and older adults

**DOI:** 10.1101/2025.05.14.653957

**Authors:** Sabrina Sghirripa, Alannah Graziano, Mitchell Goldsworthy

## Abstract

Working memory (WM) decline in ageing may be related to increases in “neural noise”, potentially reflected in the EEG aperiodic exponent. We reanalysed previously published data to investigate age-related differences in the aperiodic exponent during verbal WM and its relationship with neural inefficiency. EEG was recorded from 24 younger (18-35 years) and 30 older adults (50-86 years) during a modified Sternberg task with 1-letter, 3-letter, and 5-letter load conditions. Younger adults consistently demonstrated steeper aperiodic slopes than older adults, though this difference was less pronounced in frontal regions during retention. Unexpectedly, both age groups showed decreased (i.e. flattened) aperiodic exponents during retention relative to fixation, with minimal load-dependent effects. Notably, the relationship between task-related exponent changes and WM performance was complex and dependent on the exponent at fixation, particularly in older adults. Finally, flatter exponents during fixation and late retention were associated with greater neural inefficiency during stimulus processing, reflected by increased P3b amplitudes without corresponding WM performance improvements. These findings suggest that flatter exponents are associated with less efficient neural processing and that older adults flexibly modulate their aperiodic exponent during retention to support WM performance.

## 1 Introduction

Verbal working memory (WM) is the ability to briefly maintain and manipulate verbal information in mind to guide goal-directed behaviour and is necessary for a range of complex cognitive tasks, including language comprehension, learning, and reasoning (Baddeley, 1992). Ageing is associated with verbal WM decline, with older adults demonstrating reduced ability to hold and manipulate information in WM (Salthouse, 1994) and increased susceptibility to interference and distraction compared to younger adults (Gazzaley & D’esposito, 2007; Hasher, 2015). Age-related declines in WM can greatly impact activities of daily living and quality of later life, however, the neural processes responsible for these differences are not well understood.

Electroencephalography (EEG) studies have demonstrated that neural oscillatory activity during verbal WM changes with age (McEvoy et al., 2001; Proskovec et al., 2016; Sghirripa et al., 2021; Springer et al., 2023). However, the EEG power spectrum also contains a non-oscillatory aperiodic component, characterised by a 1/f-like decay in spectral power with increasing frequency. While traditionally dismissed as non-functional noise, aperiodic EEG activity is now understood to reflect synaptic excitation/inhibition (E/I) balance (Gao et al., 2017; Waschke et al., 2021) and neural population firing statistics (Freeman & Zhai, 2009; Manning et al., 2009; Voytek & Knight, 2015), and it has been shown to change dynamically with behavioural state (Lendner et al., 2020; Podvalny et al., 2015; Waschke et al., 2021). Notably, the aperiodic exponent—equivalent to the negative linear slope of spectral power decay in log-log space—has been reported to decrease (i.e., become flatter) in older age (McKeown et al., 2025; Merkin et al., 2023; Tran et al., 2020; Voytek et al., 2015; Waschke et al., 2017), possibly reflecting a shift toward noisier neural population spiking activity that degrades neural communication and disrupts cognitive performance (Tran et al., 2020; Voytek et al., 2015). Indeed, aperiodic activity is related to individual differences in cognitive performance in older adults (Finley et al., 2024; McKeown et al., 2025; Smith et al., 2023; Thuwal et al., 2021), with flatter spectral slopes mediating age-related declines in WM performance (Voytek et al., 2015).

There is a growing body of work examining event-induced changes in aperiodic activity during cognitive task performance. The aperiodic exponent increases (i.e., becomes steeper) after auditory (Gyurkovics et al., 2022) and visual stimulation (Manyukhina et al., 2024), in response to incongruent compared to congruent conditions of a flanker task (Jia et al., 2024), during the early foreperiod of a cued flanker task (Kałamała et al., 2024), and during “persistence-heavy” trials of a Simon Go/NoGo task (Zhang et al., 2023). For tasks engaging WM, studies have found aperiodic steepening in the peristimulus period of verbal and visuospatial n-back WM tasks (Akbarian et al., 2024; Frelih et al., 2024) and during the retention period of a continuous partial-report visual WM task (Virtue-Griffiths et al., 2022), which were related to better task performance. These findings can be interpreted within an E/I balance and neural noise framework, with task-related steepening of aperiodic exponents reflecting increased inhibitory (or decreased excitatory) processes that improve signal-to-noise ratio, allowing more precise stimulus-specific processing and higher fidelity encoding and maintenance of information in mind. Given the flattening of aperiodic exponents typically seen in older adults, a failure to steepen aperiodic activity during key stages of cognitive processing could explain age-related declines WM performance. However, whether age-related differences in task-related modulation of aperiodic activity during WM exist has yet to be seen.

Older and younger adults can also differ in how efficiently they recruit neural resources during WM and other cognitive tasks to support performance (Grady, 2012; Prince et al., 2024). While age-related over-recruitment of brain regions during cognitive tasks can sometimes reflect the engagement of compensatory processes (Cappell et al., 2010; Reuter-Lorenz & Cappell, 2008), it can also be a sign of less efficient use of neural resources, particularly in cases where increased brain activity is not matched by better behavioural performance (de Chastelaine et al., 2011; Stevens et al., 2008; Zarahn et al., 2007). A similar notion of neural efficiency has been used to explain individual differences in measures of intelligence (Neubauer & Fink, 2009). It is also one of several neural mechanisms thought to explain individual differences in resilience to brain pathology in ageing (Barulli & Stern, 2013; Stern, 2009), with higher cognitive reserve associated with higher neural efficiency measured using fMRI (Solé-Padullés et al., 2009) and EEG event-related potentials (ERPs) (Speer & Soldan, 2015). If age-related flattening of aperiodic exponents is reflective of increased neural noise, then one might expect flatter exponents during preparatory (i.e., pre-stimulus) periods of a WM task to be associated with less organised, and perhaps less efficient, neural processing.

The aims of this study were thus two-fold and were pre-registered on OSF (https://osf.io/my53h). First, we investigated age-related differences in task-related modulation of the aperiodic exponent during the retention period of a verbal WM task. Second, we examined the relationship between pre-stimulus aperiodic exponents and neural efficiency during WM encoding and probe phases, as indexed using the centroparietal P3b ERP amplitude, which is understood to reflect cognitive resource allocation (Kok, 2001; Polich, 2007) and has been used previously to study the relationship between neural efficiency and cognitive reserve (Speer & Soldan, 2015). We re-analysed data collected as part of our previous study examining age-related differences in EEG alpha frequency and power during a modified Sternberg task with 1-letter, 3-letter, and 5-letter load conditions (Sghirripa et al., 2021) to test the following hypotheses. First, the aperiodic exponent will increase (i.e., steepen) during the retention period of the task. Second, steepening during retention will be greater for younger compared to older adults and at higher compared to lower loads. Third, flatter aperiodic exponents during pre-stimulus fixation and late retention periods will be associated with reduced neural efficiency during the encoding and probe phases of the task, respectively.

## 2 Method

### 2.1 Participants

24 younger adults (mean age: 23.3 years, *SD*: 4.60, range: 18-35 years, 8 male) and 30 older adults (mean age: 62.7 years, SD: 9.09, range: 50–86 years, 17 male) participated in the study. The samples in each group were similar for years of education (older adults: *M* = 15.87 years, *SD* = 4.45 years; younger adults: *M* = 15.71 years, *SD* = 1.97). All older adults were without cognitive impairment (Addenbrooke’s Cognitive Examination score (ACE-III) > 82) (Mioshi et al., 2006). Exclusion criteria were a history of neurological or psychiatric disease, use of central nervous system altering medications, history of alcohol/substance abuse, uncorrected hearing/visual impairment, and an ACE-III score of less than 82. All participants gave informed written consent before the commencement of the study, and the experiment was approved by the University of Adelaide Human Research Ethics Committee (H-2019-154).

### 2.2 Working Memory Task

As described in Sghirripa et al. (2021), the modified Sternberg WM task used stimuli presented by PsychoPy software (Peirce, 2007) (Figure 1A). At the beginning of each trial, the participant fixated on a cross in the centre of the screen for 2 s. A memory set consisting of either 1, 3, or 5 consonants was then shown for 1 s, followed by a 4 s retention period. For load-1 and load-3 trials, the consonants were presented centrally, with filler symbols (#’s) added to maintain equal sensory input for each condition. A probe letter was then shown, and the subject was instructed to press the right arrow key on a standard keyboard if the letter was in the memory set, or the left arrow key if it was not. The probe remained on the screen until the subject responded. Probe letters were present in the memory set at 50% probability. Participants received a practice block of 20 trials to familiarise themselves with the task, before performing 20 blocks of 15 trials, yielding 300 trials overall (i.e., 100 trials per load). Each block contained an equal number of trials for each load, presented pseudorandomly, and a short break was allowed between blocks.

**Figure 1.**
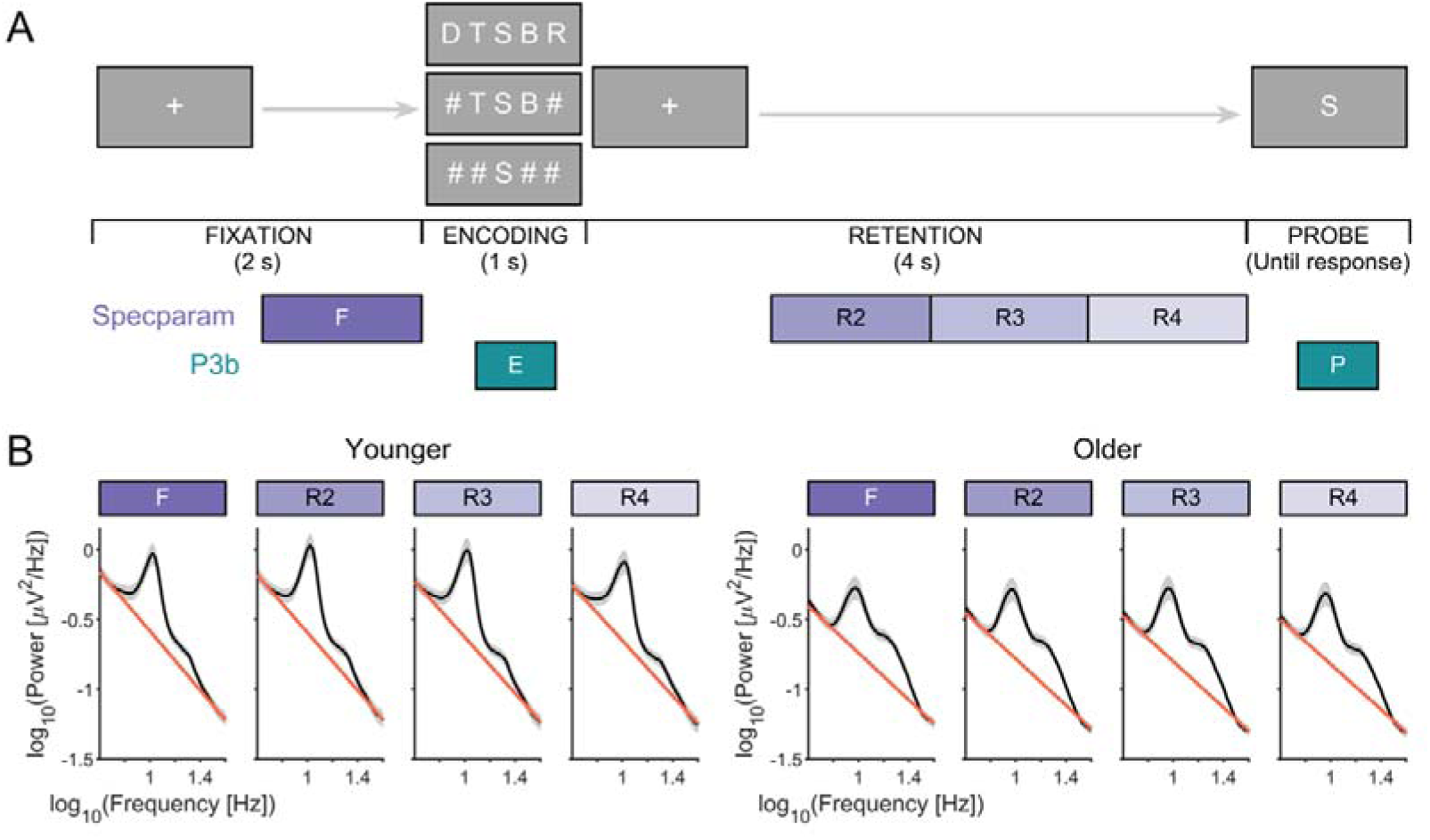
(A) Modified Sternberg task. Each trial contained four stages, including fixation lasting for 2 s; encoding, where a 1, 3, or 5 load memory set was displayed for 1 s; a 4 s retention stage and a retrieval stage where the subject responded to whether the probe was part of the memory set. Purple boxes indicate 1 s analysis periods for *specparam* (F, R2, R3, R4), while teal boxes indicate analysis period for the encoding P3b (E; 300-800 ms post memory set) and probe P3b (P; 300-800 ms post probe). (B) Log transformed power spectral density plots (black line) and aperiodic fits (red line) for younger (left) and older adults (right) at each of the analysis periods, averaged across all electrodes and WM loads.

As an additional behavioural measure not reported in the original manuscript, we calculated sensitivity (dL), which is a measure of accuracy that accounts for response bias (Snodgrass & Corwin, 1988). dL represents the difference between standardised hits and standardised false alarms and is functionally equivalent to d-prime.

### 2.3 EEG Data Acquisition

EEG data were recorded from a 64-channel cap containing Ag/AgCl scalp electrodes arranged in a 10–10 layout (Waveguard, ANT Neuro, Enschede, The Netherlands) using a Polybench TMSi EEG system (Twente Medical Systems International B.V, Oldenzaal, The Netherlands). Due to technical issues, data from the mastoids were not able to be recorded, and as such, data were recorded from 62 channels. Conductive gel was inserted into each electrode using a blunt-needle syringe to reduce impedance to < 5 kΩ. The ground electrode was located at AFz. Signals were amplified 20x, online filtered (DC-553 Hz) and sampled at 2048 Hz. Due to the lack of data from the mastoids, data were referenced to the average of all electrodes. EEG was recorded during each block of 15 trials of the WM task.

### 2.4 Data Pre-Processing

Task EEG data were pre-processed using EEGLAB (Delorme & Makeig, 2004) custom scripts using MATLAB (R2022a, The Mathworks, USA). Each block of EEG data was merged and incorrect trials, as well as trials with outlier RT (defined as > 3×SD) were flagged for removal at the epoch stage.

Noisy and unused channels were then removed based on visual inspection. The data were then band-pass (0.1–100 Hz; deviating from the original 1-100 Hz to analyse ERPs) and band-stop (48–52 Hz) filtered using zero-phase fourth-order Butterworth filters, down-sampled to 256 Hz and epoched −6.5 s to 1s relative to the beginning of the probe. Only correct trials were included in further analysis. Independent component analysis (ICA) was then conducted using the FastICA algorithm (Oja & Hyvarinen, 2000), with the “symmetric approach” and “tanh” contrast function to remove artifacts resulting from eye-blinks and persistent scalp muscle activity. Data were then checked for remaining artifact via visual inspection and trials were removed if necessary (e.g., remaining blinks, non-stereotypic artifacts). Remaining trials were then split according to memory load condition.

### 2.5 Spectral Analysis

Spectral analysis of EEG data were implemented using the FieldTrip toolbox (Oostenveld et al., 2011). EEG trials were divided into 1 s time windows, including a baseline time window prior to the memory set and multiple time windows during the retention period (Figure 1A). The first 1 s of the retention period was excluded to avoid spectral contributions from memory set ERPs. Single-trial power spectra were calculated for each time window and channel using single Hanning tapers (1-Hz frequency resolution), then averaged across trials for each memory load condition.

Parameterisation of power spectra into aperiodic and periodic components was performed using the *specparam* algorithm (Donoghue et al., 2020), implemented in FieldTrip with the following initial settings: peak width limits =D[2,12], maximum number of peaks =D6, peak threshold =D2 SD, aperiodic mode =Dfixed, and a frequency range of 2–40 Hz. Goodness-of-fit metrics (*R^2^*, mean absolute error, and frequency-wise absolute error) and visual inspection were used to determine the quality of model fits. Following the initial *specparam* fitting, goodness-of-fit metrics and visual inspection indicated poor model fit quality for certain channels. To improve fit quality, each 1 s time segment was zero-padded to 2 s, giving a 0.5-Hz frequency resolution for power spectra. *Specparam* was then repeated with the same initial settings, except with peak width limits =D[1,12].

Additionally, we identified and removed outlier channels from individual time segments that resulted in poor *specparam* model fits, which we defined as channels with an *R^2^* more than 1.5 times the interquartile range below the first quartile. Spherical spline interpolation of outlier channels was performed on time segments using *ft_channelrepair* in FieldTrip, followed by spectral analysis as described previously.

### 2.6 P3b Analysis and Neural Inefficiency Measure

A neural inefficiency measure based on Speer and Soldan (2015) was calculated using both behavioural and ERP data, which aimed to examine the relationship between behavioural performance and neural resource allocation across WM loads.

The amplitude of the P3b component for each WM load was calculated as the amplitude at the maximum peak in the 300-800 ms time window at the Pz electrode post memory set or probe. ERPs were baseline corrected to the −200-0 ms pre-stimulus period.

To calculate the neural inefficiency score, linear regression models were fitted for each participant to determine the slopes of three relationships: 1) response time (RT) as a function of WM load, 2) dL as a function of WM load, and 3) P3b peak amplitude as a function of WM load. P3b slopes were calculated separately for both encoding and probe phases. The neural inefficiency score was then calculated using the following formula:

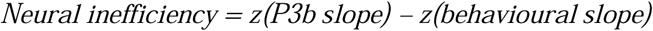

Where *z(P3b slope)* is the standardised slope of the P3b amplitude change with WM load, and *z(behavioural slope)* is either *z(dL slope)* or *-1 * z(RT slope)* (inverted so that higher values indicate better performance).

In this measure, if P3b amplitude and behavioural performance are both high, the neural inefficiency score will be lower, indicating efficient processing. Conversely, if performance is poor despite the increase in P3b amplitude, their neural inefficiency score will be higher, indicating that despite the increase in neural resources allocated to the task, there is less benefit to behavioural performance.

### 2.7 Statistical Analysis

Statistical analyses were performed using *R* version 4.4.1. Bayesian linear mixed effects models were used to analyse both behavioural and EEG data, implemented using *rstanarm* (version 2.32.6) (Gabry et al., 2024) and *bayestestR* (Makowski et al., 2019). Data visualisations were created using *ggplot2* (Wickham, Hadley, 2016) and MATLAB. Data are presented as median ± 95% HDI in figures and median [95% HDI] in text, unless indicated otherwise.

The Markov Chain Monte Carlo (MCMC) method was used to estimate the posterior distribution, and all models were run using the default weakly informative priors in the *rstanarm* package. To ensure convergence, a total of 80000 iterations on 4 chains were run with 40000 discarded for burn-in in each chain, yielding 40000 posterior samples per chain. Convergence was assessed using the potential scale reduction factor (Rhat), with values < 1.01 indicating successful convergence (Vehtari et al., 2021). Post-hoc comparisons were investigated using *emmeans* (version 1.10.3) (Lenth et al., 2025), and all 95% credible intervals were calculated using the highest density interval (HDI) method.

After running the models and confirming convergence, all inferences were drawn from the posterior distributions of the model parameters. We prioritised continuous estimation over binary hypothesis testing, reporting posterior medians and 95% HDIs to quantify effect sizes and uncertainty. To provide further context, we report two complementary measures of evidence strength that allowed us to separately evaluate whether 1) the effects exist in a certain direction, and 2) whether they are large enough to be meaningful.

First, the probability of direction (*pd*) quantifies the certainty that an effect exists in a particular direction (i.e. positive or negative). It is calculated as the proportion of the posterior distribution that shares the sign of the median, and ranges from 50% (no directional evidence) to 100% (directional certainty) (Makowski, Ben-Shachar, & Lüdecke, 2019). The following reference values for interpreting the strength of effect existence were used: *pd* > 97.5%, a likely effect; *pd* > 99%, a probable effect; and *pd* > 99.9%, a certain effect (Makowski, Ben-Shachar, Chen, et al., 2019). While *pd* correlates with frequentist *p*-values (*pd* = 97.5% ≍ *p* = 0.05), it does not depend on null hypothesis testing, instead directly quantifying the probability of an effect’s direction in the observed data (Makowski, Ben-Shachar, Chen, et al., 2019).

Second, we evaluated the practical relevance of effects by examining whether their magnitude exceeded a minimal meaningful threshold. We defined a region of practical equivalence (ROPE) as ±0.1*SD of the outcome variable, representing a threshold for trivial effect sizes (Kruschke, 2014) (i.e. negligble effect size in behavioural sciences according to (Cohen, 1988)). The practical relevance of the effects were then classified based on the proportion of their 95% HDI overlapping this ROPE. Effects with less than 1% of their HDI within the ROPE were considered practically relevant, while those with less than 2.5% overlap were considered probably relevant. Conversely, effects with more than 97.5% of their HDI overlapping the ROPE were considered probably irrelevant, and those exceeding 99% overlap were deemed irrelevant (Makowski, Ben-Shachar, & Lüdecke, 2019). Cases where 2.5-97.5% of the HDI fell within the ROPE were classified as inconclusive, indicating that the data neither strongly supported nor refuted the effect’s practical relevance. Importantly, an inconclusive result reflects uncertainty about the effect’s magnitude rather than evidence for its absence. For example, an effect might show strong evidence for existence (*pd* > 99%) while remaining inconclusive in whether the magnitude of the effect is large enough to be meaningfully important (e.g., 15% in ROPE).

To address Aim 1, we ran two Bayesian linear mixed effects models. For the first model, age group (younger vs. older), WM load (1, 3 or 5) and time segment (1 s during fixation and 1 s increments during retention period) were fixed effects, the aperiodic exponent was the outcome variable, and participant and electrode were random effects. For the second model, scalp region (frontal, central or parieto-occipital) was added as a fixed effect to determine whether the aperiodic exponent differs across regions of interest (ROIs), defined a-priori according to the 10-10 layout of the EEG cap (frontal: FP1, FPz, FP2, F7, F3, Fz, F4, F8, AF7, AF3, AF4, AF8, F5, F1, F2, F6, FT7, FT8; central: FC5, FC1, FC2, FC6, T7, C3, Cz, C4, T8, CP5, CP1, CP2, CP6, FC3, FCz, FC4, C5, C1, C2, C6, CP3, CPz, CP4, TP7, TP8; and parieto-occipital: P7, P3, Pz, P4, P8, POz, O1, Oz, O2, P5, P1, P2, P6, PO5, PO3, PO4, PO6, PO7, PO8).

As an exploratory analysis, we examined the relationship between WM performance (dL) and changes in the aperiodic exponent during retention (relative to fixation) across age groups and WM loads, controlling for retention segment. To account for effects where changes in aperiodic activity may not influence performance in a linear manner, we included both linear and nonlinear (quadratic) terms in our model.

To address Aim 2, we first ran models on the behavioural data where WM performance (RT or dL) was the outcome variable, age group and WM load were fixed effects, and participant was the random effect. Then, to determine whether there were age and WM load effects on the P3b ERP, models were run for each of the encoding and probe with P3b amplitude as the outcome, age group and WM load as fixed effects, and participant as the random effect.

To determine whether the neural inefficiency measure was related to the pre-stimulus aperiodic exponent, generalised linear models (GLM) were run for each of the encoding and probe phases with neural inefficiency as the outcome and pre-stimulus (fixation or R4) aperiodic exponent and age group as predictors. To determine whether the relationship between neural inefficiency and the pre-stimulus aperiodic exponent differed across scalp region, the same GLM was repeated using the exponent derived from either the frontal, central or parieto-occipital region.

## 3 Results

### 3.1 Aperiodic Exponent

#### 3.1.1 Age, Load and Task Effects

Across all conditions, younger adults demonstrated a larger exponent (i.e. steeper slope) than older adults (median difference = 0.21 [0.09, 0.31], *pd* = 100%, 0% in ROPE) (Figure 1B). Both age groups showed task-related changes in the exponent, with progressive decreases during retention relative to fixation (R2: −0.03 [−0.05, −0.02]; R3: −0.05 [−0.06, −0.03]; R4: −0.04 [−0.06, −0.03], all *pd* = 100%), though practical relevance was inconclusive across all segments (> 6.00% in ROPE).

WM load had minimal impact on the exponent during the task. While there was a likely effect of decreased exponents at load-5 compared to load-1 (−0.01 [−0.02, 0.00], *pd* = 97.54%), this was not practically relevant (100% in ROPE). However, the model revealed likely evidence that younger adults showed a different pattern of exponent changes at load-5 during late retention (R4) compared to older adults (0.03 [0.01, 0.06], *pd* = 98.80%, 63.24% in ROPE).

Post-hoc analyses revealed that the temporal evolution of retention-related changes differed between age groups (Figure 2A-C). At load-1, both age groups showed decreases from fixation during all retention segments (all *pd* =100%), but this was only practically relevant for younger adults in later retention periods (Younger R3/R4: 0% in ROPE). Within the retention period, while younger adults showed evidence of continued decreases from R2 to later segments (both *pd* = 100%, < 1.97% in ROPE), older adults showed little evidence for meaningful changes between retention segments (all *pd* < 96.00%, 100% in ROPE) (Figure 2A).

**Figure 2.**
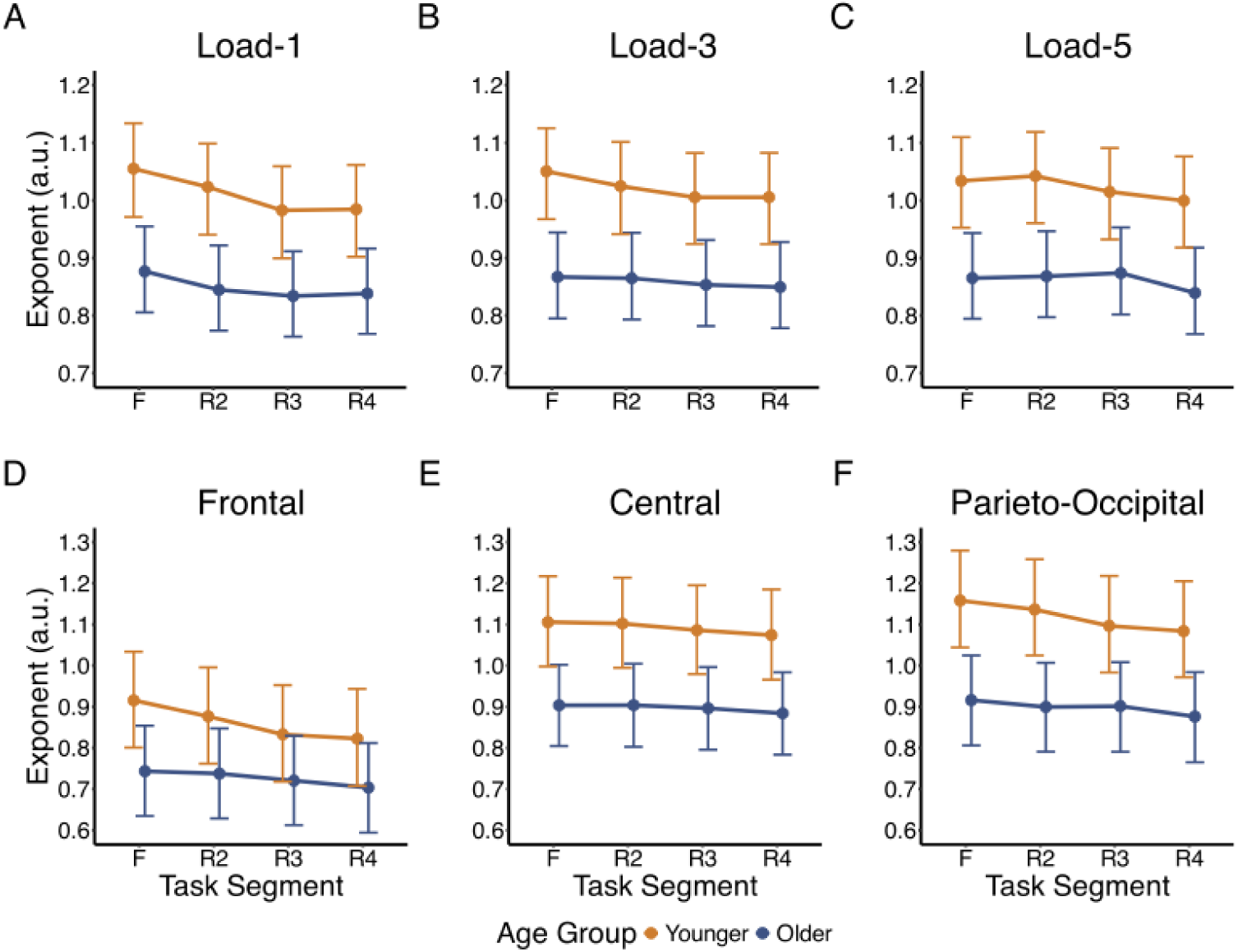
Median aperiodic exponent during task segments (F – fixation, R – retention) for younger (orange) and older (navy) adults across WM loads and ROIs. For (A), (B) and (C), the exponent is the average of all ROIs, while (D), (E) and (F) are averaged over WM load. Error bars represent 95% credible intervals based on the highest density interval (HDI).

At load-3, younger adults showed consistent decreases from fixation throughout retention, although practical relevance was mixed (all *pd* = 100%, R2: 75.42% in ROPE; R3: 1.13% in ROPE; R4: 0% in ROPE), while older adults demonstrated changes only in R3 and R4, but this was not practically relevant (R4: *pd* = 100%, both 99.11% in ROPE). Within the retention period, both age groups showed evidence for decreases from R2 to later segments (both *pd* > 99.10%) but without practical relevance (all > 82.55% in ROPE), and neither group showed changes between R3 and R4 (both *pd* < 93.60%, 100% in ROPE) (Figure 2B).

At load-5, neither age group showed early retention changes from fixation (both *pd* < 91.92%, 100% in ROPE), but both age groups demonstrated evidence for a decrease at R3/R4 with varying practical relevance (all *pd* > 98.45, > 21.21% in ROPE). Within the load-5 retention period, younger adults showed evidence for exponent decreases from R2 to later segments, but with varying practical relevance (R2 to R3: *pd* = 100%, 70.24% in ROPE; R2 to R4: *pd* = 100%, 0% in ROPE; R3 to R4: *pd* = 99.52%, 100% in ROPE). For older adults, while there was no evidence for changes in the exponent from R2 to R3 (*pd* = 63.23%), there was evidence for decreases from both R2 to R4 and R3 to R4 (both *pd* = 100%), all with inconclusive practical relevance (all > 12.34% in ROPE) (Figure 2C).

#### 3.1.2 Regional Differences

The model with ROI added as a fixed effect demonstrated differences in aperiodic exponents across regions, with frontal regions showing lower exponents compared to central (median difference = −0.16 [−0.27, −0.06], *pd* = 99.75%, 0% in ROPE), but not parieto-occipital regions (median difference = 0.01 [−0.10, 0.11], *pd* = 57%, 53% in ROPE). There was evidence for an interaction between age group and parieto-occipital ROI (median difference = 0.05 [0.02, 0.09], *pd* = 99.90%, 15.97% in ROPE), and for younger adults having distinct changes in frontal regions during later retention periods compared to older adults (R3: *pd* = 99.40%, 13.42% in ROPE; R4: *pd* = 98.98%, 16.68% in ROPE). Similar effects were observed in parieto-occipital regions during R3 (*pd* = 98.83%, 17.63% in ROPE), however, practical relevance was inconclusive in all interactions. No effects involving WM load reached the threshold for probable directional evidence or practical relevance (all pd < 97.20%, > 11.97% in ROPE).

Post-hoc comparisons examining age differences across segments and ROIs revealed that while young adults consistently showed larger exponents than older adults, the magnitude of this difference varied by ROI and task stage. Central and parieto-occipital regions showed practically relevant age differences across all segments (all *pd* > 99.95, 0% in ROPE) (Figure 2E-F). In contrast, frontal regions showed strong age differences at fixation (*pd* = 99.83%, 0% in ROPE) that diminished during late retention (R3/R4: *pd* > 96.65%, > 5.82% in ROPE; Figure 2D).

#### 3.1.3 Association with WM performance

The model examining the relationship between the change in exponent during retention (relative to fixation) and dL revealed that changes in the aperiodic exponent during retention demonstrated a U-shaped relationship with dL (median = 7.14 [1.13, 12.84], *pd* = 99.08%, 0% in ROPE). There was suggestive but inconclusive evidence for age group differences in the quadratic relationship, with the practical relevance threshold met but directional evidence falling below the threshold for a likely effect (median = −7.85 [−16.89, 0.95], *pd* = 96.27%, 0.53% in ROPE) (Figure 3A), indicating that larger exponent changes (both positive and negative) were associated with higher dL in older adults.

**Figure 3.**
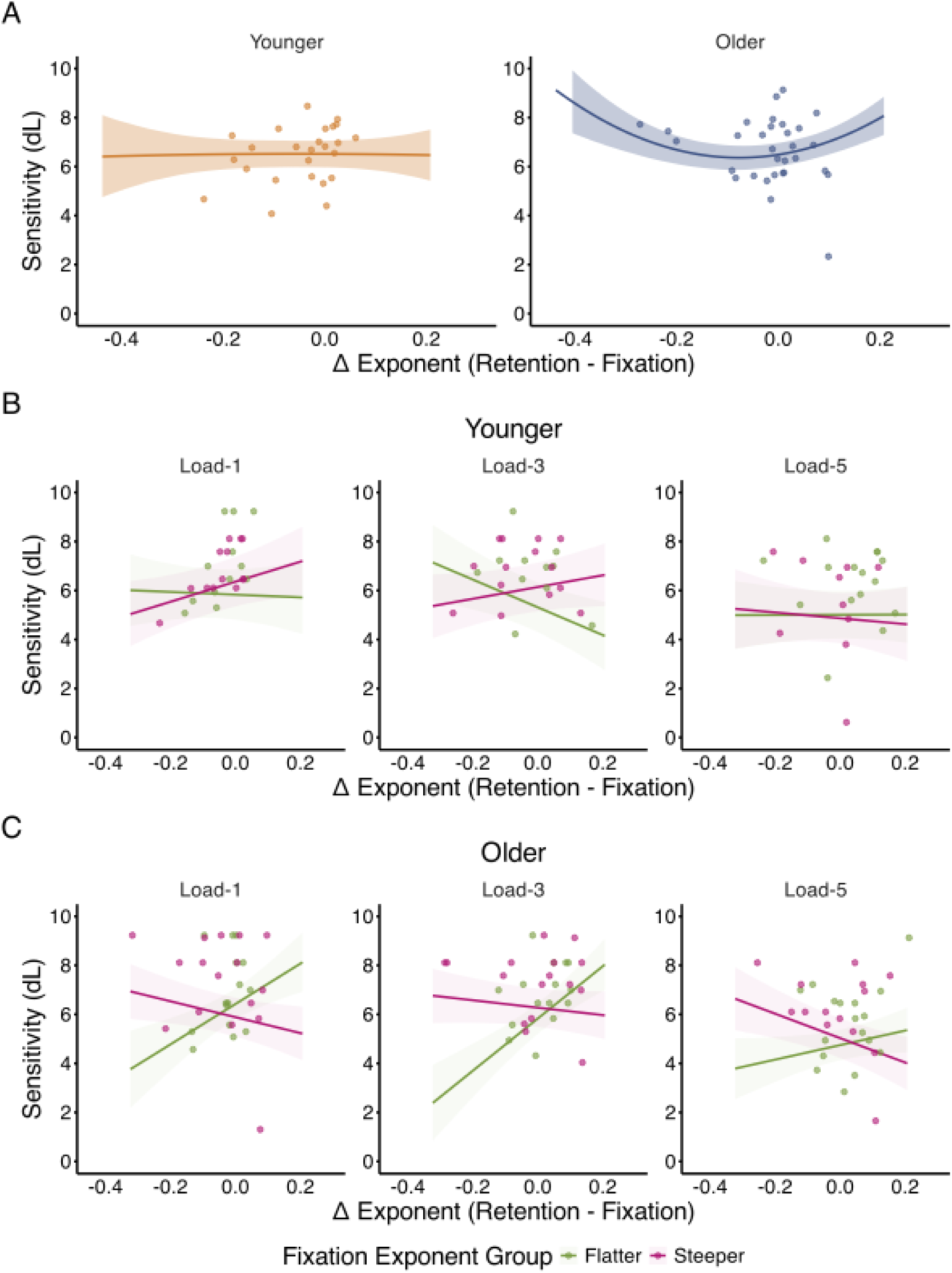
Relationship between the change in aperiodic exponent from fixation to retention period and sensitivity (dL). (A) Overall quadratic relationship between change in exponent and dL for younger (left) and older (right) adults, showing a more pronounced U-shaped relationship in older adults. Load-dependent relationships between exponent change and dL in (B) younger and (C) older adults, split by fixation exponent group (Flatter vs Steeper). Shaded areas represent 95% HDI.

To further understand how the fixation exponent influenced the relationship between exponent changes during retention and dL, we conducted a follow-up analysis that categorised participants based on their fixation exponent (i.e., steeper or flatter based on a median split). The model revealed evidence for a linear interaction between exponent change, fixation exponent group, and age group (median = 16.26 [8.43, 23.92], *pd* = 100%, 0% in ROPE), that was further modulated by WM load (load-5; median = −9.52 [−18.78, −0.07], *pd* = 97.62%, 0% in ROPE) (Figure 3B-C).

Post-hoc analyses revealed that at load-1, better WM performance in older adults was associated with retention exponents changing in the opposite direction from their fixation exponent (flatter at fixation: 0.83 [0.38, 1.30]; steeper at fixation: −0.33 [−0.62, −0.04], both *pd* > 99.36, 0% in ROPE). In younger adults, better WM performance was associated with further steepening during retention, but only for those with steeper fixation exponents (0.41 [0.07, 0.75], *pd* = 98.40%, 0% in ROPE; flatter fixation exponents: −0.05 [−0.46, 0.36], *pd* = 76.44%, 3.02% in ROPE).

At load-3, older adults with flatter fixation exponents performed better when their exponent steepened during retention (1.06 [0.64, 1.48], *pd* = 100%, 0% in ROPE), while for younger adults with flatter fixation exponents, better performance was associated with further flattening (−0.57 [−0.96, −0.17], *pd* = 99.91%, 0% in ROPE). Neither age group showed certain effects with steeper baselines (older: −0.16 [−0.41, 0.11]; younger: 0.24 [−0.09, 0.56], both *pd* < 93.46%, > 2.21% in ROPE).

At load-5, flattening during retention was associated with better performance in older adults with steeper fixation exponents (−0.50 [−0.84, −0.19], *pd* = 99.95%, 0% in ROPE), with no certain effects in other conditions (older/flatter at fixation: 0.29 [−0.05, 0.60]; younger/flatter at fixation: 0.00 [−0.31, 0.30]; younger/steeper at fixation: −0.12 [−0.56, 0.34], all *pd* < 83.48%, > 2.35% in ROPE).

### 3.2 Neural Inefficiency

#### 3.2.1 Behavioural Data

WM performance data have already been published in the original manuscript. For detailed response time (RT) and WM capacity results, see Sghirripa et al. (2021).

To characterise the additional behavioural measure for neural inefficiency calculations, we ran a Bayesian linear mixed effects model with sensitivity (dL) as the outcome. The model revealed a certain effect of WM load on dL, with performance declining in load-5 compared to load-1 (median difference = −1.18 [−1.74, −0.63], *pd* = 100%, 0% in ROPE), but not at load-3 (median difference = −0.005 [−0.60, 0.51], *pd* = 50.60%, 44.26% in ROPE). Similarly, there was little evidence for an overall age difference in dL (*pd* = 64.85%, 29.58% in ROPE), nor for an interaction between age group and load (both *pd* < 70.08%, > 28.08% in ROPE), suggesting that the impact of increasing WM load on dL was similar across age groups (Figure 4A).

**Figure 4.**
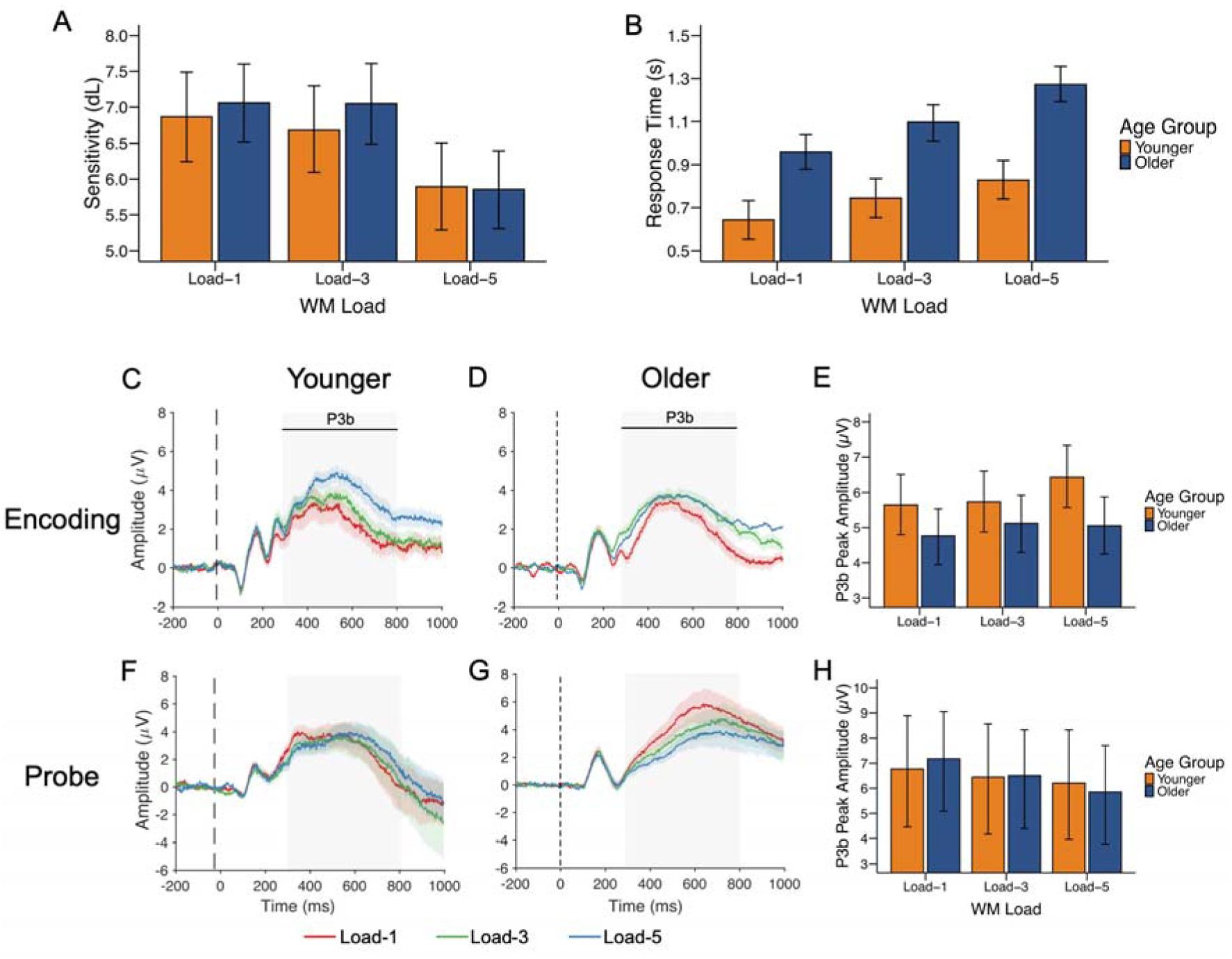
Behavioural performance and ERP responses during the WM task for younger (orange) and older (navy) adults. (A) Sensitivity (dL) and (B) response time across WM loads. (C-E) ERP responses during encoding: time course for younger (C) and older (D) adults with corresponding P3b peak amplitudes (E). (F-H) ERP responses during probe presentation: time course for younger (F) and older (G) adults with corresponding P3b peak amplitudes (H). Shaded areas in ERP plots indicate the P3b time window used for peak amplitude calculation. Error bars represent 95% credible intervals based on the highest density interval (HDI).

#### 3.2.2 P3b Components

For the peak amplitude of the P3b component during encoding, there were uncertain effects of age (*pd* = 94.42%, 9.95% in ROPE) and WM load (load-3/load-5: *pd* < 88.60%, > 30.61% in ROPE), and little evidence for an interaction between age-group and WM load (load-3:/load-5 *pd* < 87.58%, > 22.82% in ROPE) (Figure 4C-D).

Given our focus on neural efficiency, to better understand age- and load-related modulation of the P3b component, post-hoc comparisons were conducted. These revealed that younger adults showed larger P3b amplitudes compared to older adults at load-5 (*pd* = 99.20%, 0% in ROPE) and demonstrated increased amplitude from load-1 to load-5 (*pd* = 99.55%, 0.87% in ROPE), while older adults showed little evidence of amplitude modulation between these loads (*pd* = 84.17%, 38.37% in ROPE) (Figure 4E).

At the probe stage, there was a certain and practically relevant decrease in P3b amplitude at load-5 compared with load-1 (median difference = −1.30 µV [−1.98, −0.60], *pd* = 100%, 0% in ROPE). This effect was particularly pronounced in older adults (median difference = 1.31 µV [0.63, 2.01], pd = 99.98%, 0% in ROPE) (Figure 4G-H) while younger adults showed no strong evidence for load-dependent modulation (*pd* = 91.47%, 49.08% in ROPE) (Figure 4F-H).

#### 3.2.3 Encoding

There was evidence for a relationship between neural inefficiency calculated from the encoding P3b and dL, and the aperiodic exponent during fixation, with smaller exponents (i.e. flatter slopes) associated with greater neural inefficiency (median = −2.92 [−5.52, −0.31], *pd* = 98.32%, 0% in ROPE) (Figure 5A). However, there was no clear evidence for age-related differences in neural inefficiency (median difference = −0.94 [−4.69, 2.74], *pd* = 69.14%, 6.28% in ROPE), or an interaction between age group and fixation exponent (median = 1.35 [−2.27, 5.14], *pd* = 76.48%, 5.48% in ROPE).

**Figure 5.**
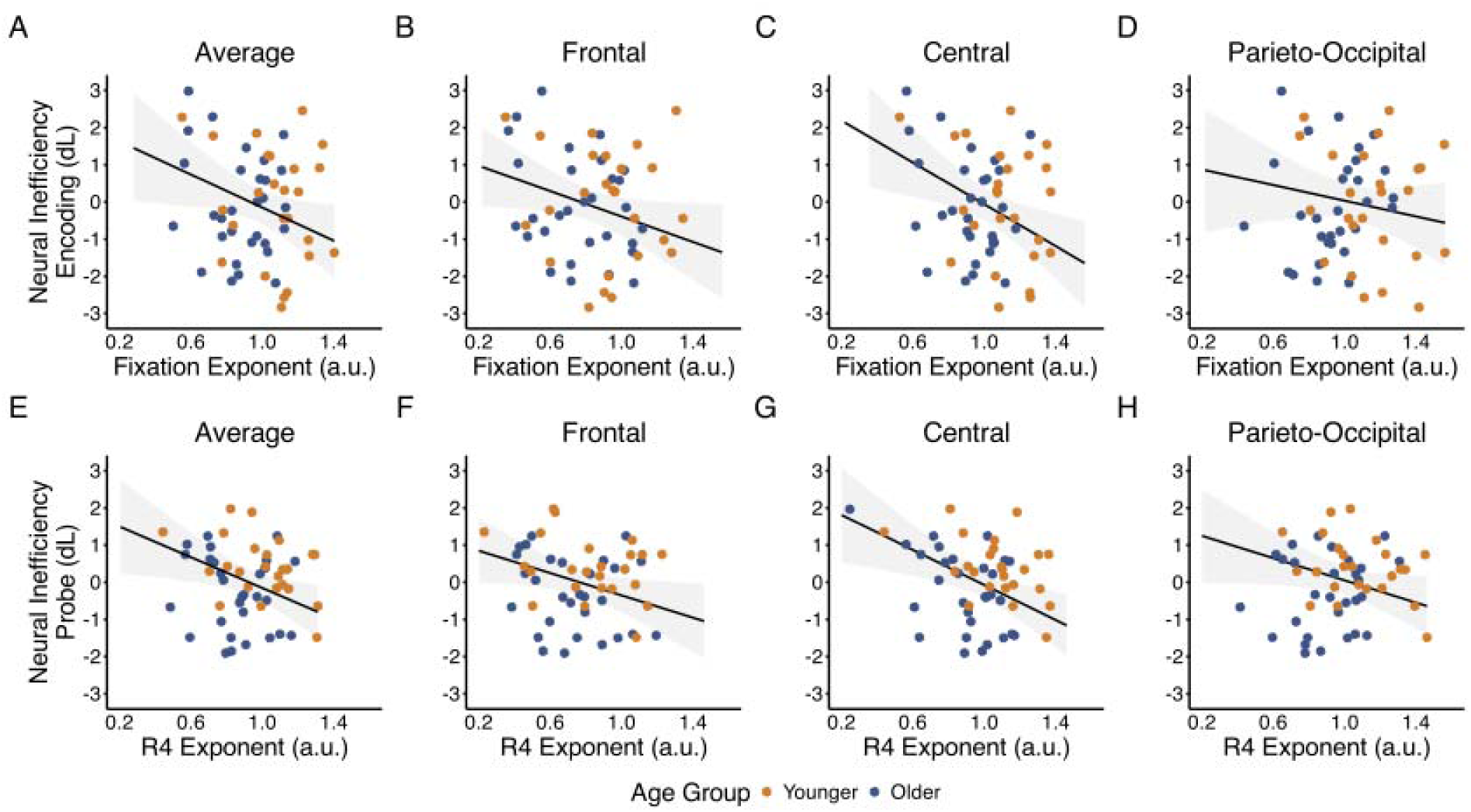
Relationship between pre-stimulus aperiodic exponents and neural inefficiency across age groups and ROIs. Neural inefficiency scores calculated during encoding (A-D) and probe phases (E-H) are plotted against aperiodic exponents measured during fixation (top row) and the final retention period R4 (bottom row), respectively. Each column represents the following ROIs: average across all electrodes (A, E), frontal (B, F), central (C, G), and parieto-occipital regions (D, H). Orange dots represent younger adults, and blue dots represent older adults. Black lines show the estimated linear fit across all participants, and shaded grey represents the 95% HDI for the fit.

To understand regional effects, separate models were run using the fixation exponent derived from each ROI. The models revealed robust evidence for a relationship between greater neural inefficiency with smaller (i.e. flatter) fixation exponents in both frontal (median = −2.85 [−4.93, −0.60], *pd* = 99.60%, 0% in ROPE) and central regions (median = −3.50 [−6.09, −1.03], *pd* = 99.50%, 0% in ROPE), but not in the parieto-occipital region (median = −1.19 [−3.58, 1.33], *pd* = 82.55%, 6.47% in ROPE) (Figure 5B-D). In all models, there was little evidence for an interaction between age group and the fixation exponent on neural inefficiency (all *pd* < 92.53, all > 2.84% in ROPE).

Another model examining RT-based neural inefficiency showed little evidence for relationships with the fixation exponent (median = −1.80 [−4.21, 0.64], *pd* = 92.64%, 3.41% in ROPE), age group (*pd* = 52.73%, 7.03% in ROPE), or an interaction (*pd* = 54.84, 7.20% in ROPE). ROI-specific models revealed that smaller frontal fixation exponents were associated with greater RT neural inefficiency (median = −2.18 [−4.31, −0.25], *pd* = 98.58%, 0% in ROPE), but showed no evidence for effects or interactions in other regions (all *pd* < 94.47%, > 2.93% in ROPE).

#### 3.2.4 Probe

There was also evidence for a relationship between the exponent during the final retention segment (R4) and neural inefficiency in response to the probe (median = −2.03 [−4.01, 0.02], *pd* = 97.63%, 0.61% in ROPE) (Figure 5E). There was little evidence for an effect of age group (median = 0.70 [−2.07, 3.60], *pd* = 68.84%, 6.63% in ROPE) or an interaction between age group and R4 exponent on neural inefficiency (median = −0.06 [−3.06, 2.90], *pd* = 51.73%, 7.27% in ROPE).

ROI based models demonstrated evidence for a relationship between R4 exponent in the central region and neural inefficiency to the probe, with smaller exponents associated with greater neural inefficiency (median = −2.84 [−4.78, −0.71], *pd* = 99.48%, 0% in ROPE) (Figure 5G). For the frontal region, there was a trend towards a relationship between the R4 exponent and neural inefficiency (median = −1.71 [−3.55, 0.07], *pd* = 96.83%, 2.18% in ROPE), but this did not meet the threshold for a probable effect (Figure 5F), while no consistent effect was seen for the parieto-occipital region (median = −0.86 [−2.70, 1.06], *pd* = 81.77%, 6.92% in ROPE) (Figure 5H). In all models, there was little evidence for an interaction between age group and R4 exponent on probe neural inefficiency (all pd < 71.50%, all > 4.84% in ROPE).

The model using the RT neural inefficiency score demonstrated a trend towards a relationship between the R4 exponent and neural inefficiency (median = −2.22 [−4.87, 0.28], *pd* = 95.25%, 2.29% in ROPE), but no evidence for an age group effect (median = −1.06 [−4.29, 2.30], *pd* = 73.67%, 4.89% in ROPE) and no evidence for an interaction between age group and the R4 exponent (median = 0.91 [−2.52, 4.34], *pd* = 69.92%, 5.03% in ROPE). Although directional evidence trended towards a relationship between the R4 exponent and neural inefficiency in all ROIs, these did not meet the threshold for a probable effect (all *pd* <95.47%, > 2.26% in ROPE).

## 4 Discussion

In this study, we investigated age-related differences in the aperiodic exponent during a verbal WM task, and the relationship between the aperiodic exponent and neural inefficiency. We found that: 1) unexpectedly, the aperiodic exponent was reduced (i.e. the slope flattened) during the retention period, but was unaffected by WM load; 2) younger adults had larger aperiodic exponents (i.e. steeper aperiodic slopes) than older adults; 3) changes in the aperiodic exponent during retention showed load-dependent relationships with performance that varied based on the exponent at fixation, particularly in older adults; and 4) flatter exponents at fixation and before the probe were associated with greater neural inefficiency, particularly in frontal and central regions.

### 4.1. Relative to fixation, the aperiodic exponent flattens during WM retention, irrespective of WM load

Inconsistent with our hypothesis, a decrease in the aperiodic exponent during the retention period relative to fixation was observed, with this change occurring across WM loads and age groups. The direction in which the aperiodic exponent is modulated during cognitive task performance shows mixed patterns across studies, with some reporting steepening while others report flattening of the slope during tasks. Steepening of the exponent has been reported in studies using a cued flanker task (Kałamała et al., 2024), auditory stimulation tasks (Gyurkovics et al., 2022), verbal WM (Frelih et al., 2024) and visual WM tasks (Virtue-Griffiths et al., 2022). However, the flattening of the aperiodic exponent during retention observed here is consistent with work detailing the pattern of aperiodic activity in visual WM tasks (Donoghue et al., 2020), and aligns with ECoG studies showing decreased 1/f exponents across visual and auditory stimuli (Podvalny et al., 2015). Similar results have been seen in EEG recordings, in which occipital exponents decreased under visual compared to auditory attention, providing evidence for a modality-specific flattening of EEG aperiodic slopes through the selective allocation of attentional resources (Waschke et al., 2021).

There is no clear consensus on the physiological basis of task-related modulations of aperiodic exponents observed in these previous studies, leading to several possible explanations for the flattening during WM retention we observed. Evidence suggests that a flatter exponent reflects an increase in background asynchronous activity (i.e. neural noise) (Freeman & Zhai, 2009; Manning et al., 2009), which manifests at the network level as a shift towards excitation relative to inhibition (i.e. an increased E/I ratio) (Gao et al., 2017). During WM retention, these network-level changes could be functionally advantageous, particularly in frontal regions, in which the increased neural excitability and asynchronous firing could support the sustained neural activity required to maintain information in WM without ongoing sensory input during retention (Goldman-Rakic, 1995; Lim & Goldman, 2013).

High-frequency activity bursts during WM retention offer another potential mechanism (Lundqvist et al., 2016). Perhaps the flattening of the aperiodic slope may capture the population-level dynamics of these bursts, which transiently elevate neural firing rates and increase broadband high-frequency activity. This interpretation aligns with Hong and Rebec’s (2012) compensatory framework, where brief firing rate increases could efficiently maintain information through periods of temporarily increased excitation and desynchronised activity, which might be captured in the flattening of the aperiodic exponent (Hong & Rebec, 2012). However, given the uncertain practical relevance of many of our results, further research is needed to understand the functional relevance of aperiodic slope flattening during WM retention.

### 4.2. The relationship between task-related aperiodic exponents and WM performance differs by age group

The relationship between age and the exponent during the task revealed several key insights about the role of the aperiodic exponent in WM performance. Consistent with a growing body of literature (Dave et al., 2018; Donoghue et al., 2020; Merkin et al., 2023; Voytek et al., 2015), we observed flatter aperiodic slopes in older adults compared to younger adults, though this difference became less pronounced in frontal regions during retention. Several mechanisms have been proposed to explain these flatter slopes in ageing, including a decrease in the reliability of neural communication resulting in ‘noisier’ neural activity (Crossman & Szafran, 1956; Voytek & Knight, 2015), shifts in synaptic excitation/inhibition balance (Gao et al., 2017) and age-related changes in brain tissue properties (Bédard & Destexhe, 2009). Notably, while these age differences persisted during our task, both groups achieved similar WM performance across loads, challenging previous associations between flatter slopes and poorer WM performance in older adults (Voytek et al., 2015).

Rather than age-related flatter exponents being universally disadvantageous for WM performance, we found that the ability to modulate the exponent during the task was associated with better WM performance, particularly in older adults. Those with flatter baseline exponents performed better when able to steepen exponents during retention at load-1 and load-3, but not at load-5, suggesting a potential compensatory mechanism that has limited capacity. In contrast, older adults with steeper baselines also benefited from flattening during retention, particularly at load-5.

These load-dependent patterns of exponent modulation may reflect distinct neural strategies for WM retention. The steepening of exponents in those with flatter baselines could represent an attempt to improve the signal-to-noise ratio and the precision of the representation of items in WM, which may help compensate for less precise encoding of the memory set (Reuter-Lorenz & Cappell, 2008; Voytek & Knight, 2015). Similarly, while the flattening observed in those with steeper baselines aligns with the typical task-related flattening we observed across conditions, its relationship with performance at load-5 suggests it might represent an adaptive neural strategy rather than just typical task-related changes, where an increase in excitatory activity may be required to maintain items in the WM store, perhaps through active verbal rehearsal.

In contrast, younger adults performed better when changes in their exponents followed their fixation patterns during retention; those with steeper baselines benefited from further steepening, while those with flatter baselines benefited from further flattening. However, these relationships were less consistent across loads. This age difference could reflect greater individual variability in how older adults successfully engage with the task. While younger adults show more uniform neural strategies aligned with their baseline states, older adults may employ different strategies depending on their baseline characteristics. For example, younger adults might rely on active verbal rehearsal, while older adults’ varied exponent modulation could reflect either verbal rehearsal or greater engagement of inhibitory control processes to maintain task-relevant information. However, these interpretations remain speculative and highlight the importance of considering both baseline characteristics and dynamic modulation when investigating how aperiodic activity relates to WM performance across the lifespan.

### 4.3. Flatter pre-stimulus aperiodic exponents are associated with neural inefficiency in both age groups

Beyond these patterns of exponent modulation, we also found that flatter slopes at fixation and prior to the probe were associated with less efficient neural processing across both age groups, with individuals showing increased neural resource recruitment, indexed by P3b amplitude, that was not matched by better behavioural performance (Speer & Soldan, 2015). If the aperiodic exponent decreases reflect an increase in neural noise due to an increase in asynchronous neural firing (Freeman & Zhai, 2009; Manning et al., 2009), leading to a decreased signal-to-noise ratio (Voytek & Knight, 2015), then perhaps a flatter exponent during preparatory periods (both at fixation and prior to the probe) represents a less organised neural state. In this noisier state, additional neural resources may be required to distinguish task-relevant signals from background noise to maintain stable WM representations. Notably, flatter exponents during late retention were also linked to neural inefficiency to the probe stimulus, suggesting that while initial exponent flattening may support WM retention, steeper exponents during extended retention could preserve better signal-to-noise ratios and enable more efficient probe processing.

Together, our findings reveal a nuanced relationship between aperiodic activity and verbal WM performance that varies with age. Despite showing consistently flatter exponents, older adults demonstrated the same link between flatter exponents and neural inefficiency as younger adults during both encoding and probe processing. During retention, however, older adults with flatter baseline exponents benefited most from modulating their aperiodic activity away from their fixation state, particularly at low and moderate loads. This pattern is consistent with the Compensation-Related Utilisation of Neural Circuits Hypothesis (CRUNCH; Reuter-Lorenz & Cappell, 2008), which suggests that the ageing brain may work harder to compensate for declining efficiency or processing deficiencies, in this case, by flexibly modulating the exponent during retention after less efficient stimulus processing.

However, the effectiveness of these compensatory mechanisms appears to break down at higher loads for older adults with flatter fixation exponents, perhaps because the combined demands of inefficient processing and compensatory modulation exceed available neural resources. Meanwhile, the more variable patterns in younger adults may suggest the task demands were insufficient to necessitate compensatory strategies, even in those with reduced neural efficiency.

### 4.4 Limitations

Our findings should be considered in light of several limitations. Notably, we found no age differences in WM performance, neural inefficiency, or remarkably different patterns of task-related exponent modulation, despite consistently flatter aperiodic exponents in older adults. Our task may have been insufficient to capture age differences in WM performance due to its relatively low loads and verbal nature, as verbal WM shows more resistance to age-related decline (Hale et al., 2011). This may also explain the absence of age-related differences in neural inefficiency (Daffner et al., 2011; Osaka et al., 2007). However, the presence of potential compensatory patterns in older adults despite equivalent performance may suggest proactive engagement of neural strategies even under low task demands. Future studies using higher WM loads or more challenging conditions may better reveal age-specific relationships between the aperiodic exponent and neural efficiency.

### 4.5. Conclusion

Here, we provide novel insights into the association between the aperiodic exponent and verbal WM in cognitive ageing, demonstrating that flatter exponents are associated with less efficient neural processing and that older adults flexibly modulate their exponent during retention to support WM performance. Future research should now focus on understanding the physiological underpinnings of the aperiodic exponent during task performance and how its modulation supports cognitive function across the lifespan.

## Funding Sources

This work was supported by the Australian Research Council (ARC) [grant number: FT230100658].

## Notes

### Competing Interest Statement

The authors have declared no competing interest.

### Summary of Updates

Author order error corrected, no other changes made.

## References

Akbarian, F., Rossi, C., Costers, L., D’hooghe, M. B., D’haeseleer, M., Nagels, G., & Van Schependom, J. (2024). Stimulus-related modulation in the 1/f spectral slope suggests an impaired inhibition during a working memory task in people with multiple sclerosis. Multiple Sclerosis Journal, 30(8), 1036–1046. 10.1177/13524585241253777

Baddeley, A. (1992). Working memory. Science, 255(5044), 556–559. 10.1126/science.1736359

Barulli, D., & Stern, Y. (2013). Efficiency, capacity, compensation, maintenance, plasticity: Emerging concepts in cognitive reserve. Trends in Cognitive Sciences, 17(10), 502–509. 10.1016/j.tics.2013.08.012

Bédard, C., & Destexhe, A. (2009). Macroscopic Models of Local Field Potentials and the Apparent 1/f Noise in Brain Activity. Biophysical Journal, 96(7), 2589–2603. 10.1016/j.bpj.2008.12.3951

Cappell, K. A., Gmeindl, L., & Reuter-Lorenz, P. A. (2010). Age differences in prefontal recruitment during verbal working memory maintenance depend on memory load. Cortex, 46(4), 462–473. 10.1016/j.cortex.2009.11.009

Cohen, J. (1988). *Statistical Power Analysis for the Behavioral Sciences* (2nd ed.). Routledge. 10.4324/9780203771587

Crossman, E. R., & Szafran, J. (1956). Changes with age in the speed of information-intake and discrimination. *Experientia*, Suppl 4, 128–134; discussion, 135.

Daffner, K. R., Chong, H., Sun, X., Tarbi, E. C., Riis, J. L., McGinnis, S. M., & Holcomb, P. J. (2011). Mechanisms Underlying Age- and Performance-related Differences in Working Memory. Journal of Cognitive Neuroscience, 23(6), 1298–1314. 10.1162/jocn.2010.21540

Dave, S., Brothers, T. A., & Swaab, T. Y. (2018). 1/*f* neural noise and electrophysiological indices of contextual prediction in aging. Brain Research, 1691, 34–43. 10.1016/j.brainres.2018.04.007

de Chastelaine, M., Wang, T. H., Minton, B., Muftuler, L. T., & Rugg, M. D. (2011). The Effects of Age, Memory Performance, and Callosal Integrity on the Neural Correlates of Successful Associative Encoding. Cerebral Cortex, 21(9), 2166–2176. 10.1093/cercor/bhq294

Donoghue, T., Haller, M., Peterson, E. J., Varma, P., Sebastian, P., Gao, R., Noto, T., Lara, A. H., Wallis, J. D., Knight, R. T., Shestyuk, A., & Voytek, B. (2020). Parameterizing neural power spectra into periodic and aperiodic components. Nature Neuroscience, 23(12), Article 12. 10.1038/s41593-020-00744-x

Finley, A. J., Angus, D. J., Knight, E. L., Reekum, C. M. van, Lachman, M. E., Davidson, R. J., & Schaefer, S. M. (2024). Resting EEG Periodic and Aperiodic Components Predict Cognitive Decline Over 10 Years. Journal of Neuroscience, 44(13). 10.1523/JNEUROSCI.1332-23.2024

Freeman, W. J., & Zhai, J. (2009). Simulated power spectral density (PSD) of background electrocorticogram (ECoG). Cognitive Neurodynamics, 3(1), 97–103. 10.1007/s11571-008-9064-y

Frelih, T., Matkovič, A., Mlinarič, T., Bon, J., & Repovš, G. (2024). Modulation of aperiodic EEG activity provides sensitive index of cognitive state changes during working memory task (p. 2024.05.13.593835). bioRxiv. 10.1101/2024.05.13.593835

Gabry, J., Ali, I., Brilleman, S., Novik (R/stan_jm.R), J. B., AstraZeneca (R/stan_jm.R), University, T. of C., Wood (R/stan_gamm4.R), S., Team (R/stan_aov.R), R. C. D., Bates (R/pp_data.R), D., Maechler (R/pp_data.R), M., Bolker (R/pp_data.R), B., Walker (R/pp_data.R), S., Ripley (R/stan_aov.R, B., R/stan_polr.R), Venables (R/stan_polr.R), W., Burkner (R/misc.R), P.-C., & Goodrich, B. (2024). rstanarm: Bayesian Applied Regression Modeling via Stan (Version 2.32.1) [Computer software]. https://cloud.r-project.org/web/packages/rstanarm/index.html

Gao, R., Peterson, E. J., & Voytek, B. (2017). Inferring synaptic excitation/inhibition balance from field potentials. NeuroImage, 158, 70–78. 10.1016/j.neuroimage.2017.06.078

Gazzaley, A., & D’esposito, M. (2007). Top-Down Modulation and Normal Aging. Annals of the New York Academy of Sciences, 1097(1), 67–83. 10.1196/annals.1379.010

Goldman-Rakic, P. S. (1995). Cellular basis of working memory. Neuron, 14(3), 477–485. 10.1016/0896-6273(95)90304-6

Grady, C. (2012). The cognitive neuroscience of ageing. Nature Reviews Neuroscience, 13(7), 491–505. 10.1038/nrn3256

Gyurkovics, M., Clements, G. M., Low, K. A., Fabiani, M., & Gratton, G. (2022). Stimulus-Induced Changes in 1/f-like Background Activity in EEG. Journal of Neuroscience, 42(37), 7144–7151. 10.1523/JNEUROSCI.0414-22.2022

Hale, S., Rose, N. S., Myerson, J., Strube, M. J., Sommers, M., Tye-Murray, N., & Spehar, B. (2011). The structure of working memory abilities across the adult life span. Psychology and Aging, 26(1), 92–110. 10.1037/a0021483

Hasher, L. (2015). Inhibitory deficit hypothesis. The Encyclopedia of Adulthood and Aging, 1–5.

Hong, S. L., & Rebec, G. V. (2012). A new perspective on behavioral inconsistency and neural noise in aging: Compensatory speeding of neural communication. Frontiers in Aging Neuroscience, 4. 10.3389/fnagi.2012.00027

Jia, S., Liu, D., Song, W., Beste, C., Colzato, L., & Hommel, B. (2024). Tracing conflict-induced cognitive-control adjustments over time using aperiodic EEG activity. Cerebral Cortex, 34(5), bhae185. 10.1093/cercor/bhae185

Kałamała, P., Gyurkovics, M., Bowie, D. C., Clements, G. M., Low, K. A., Dolcos, F., Fabiani, M., & Gratton, G. (2024). Event-induced modulation of aperiodic background EEG: Attention-dependent and age-related shifts in E:I balance, and their consequences for behavior. Imaging Neuroscience, 2, 1–18. 10.1162/imag_a_00054

Kok, A. (2001). On the utility of P3 amplitude as a measure of processing capacity. Psychophysiology, 38(3), 557–577. 10.1017/S0048577201990559

Kruschke, J. (2014). Doing Bayesian Data Analysis: A Tutorial with R, JAGS, and Stan. Academic Press.

Lendner, J. D., Helfrich, R. F., Mander, B. A., Romundstad, L., Lin, J. J., Walker, M. P., Larsson, P. G., & Knight, R. T. (2020). An electrophysiological marker of arousal level in humans. eLife, 9, e55092. 10.7554/eLife.55092

Lenth, R. V., Banfai, B., Bolker, B., Buerkner, P., Giné-Vázquez, I., Herve, M., Jung, M., Love, J., Miguez, F., Piaskowski, J., Riebl, H., & Singmann, H. (2025). emmeans: Estimated Marginal Means, aka Least-Squares Means (Version 1.11.1) [Computer software]. https://cran.r-project.org/web/packages/emmeans/index.html

Lim, S., & Goldman, M. S. (2013). Balanced cortical microcircuitry for maintaining information in working memory. Nature Neuroscience, 16(9), 1306–1314. 10.1038/nn.3492

Lundqvist, M., Rose, J., Herman, P., Brincat, S. L., Buschman, T. J., & Miller, E. K. (2016). Gamma and Beta Bursts Underlie Working Memory. Neuron, 90(1), 152–164. 10.1016/j.neuron.2016.02.028

Makowski, D., Ben-Shachar, M. S., Chen, S. H. A., & Lüdecke, D. (2019). Indices of Effect Existence and Significance in the Bayesian Framework. Frontiers in Psychology, 10. 10.3389/fpsyg.2019.02767

Makowski, D., Ben-Shachar, M. S., & Lüdecke, D. (2019). bayestestR: Describing effects and their uncertainty, existence and significance within the Bayesian framework. Journal of Open Source Software, 4(40), 1541.

Manning, J. R., Jacobs, J., Fried, I., & Kahana, M. J. (2009). Broadband Shifts in Local Field Potential Power Spectra Are Correlated with Single-Neuron Spiking in Humans. Journal of Neuroscience, 29(43), 13613–13620. 10.1523/JNEUROSCI.2041-09.2009

Manyukhina, V. O., Prokofyev, A. O., Obukhova, T. S., Stroganova, T. A., & Orekhova, E. V. (2024). Changes in high-frequency aperiodic 1/f slope and periodic activity reflect post-stimulus functional inhibition in the visual cortex. Imaging Neuroscience, 2, 1–24. 10.1162/imag_a_00146

McEvoy, L. K., Pellouchoud, E., Smith, M. E., & Gevins, A. (2001). Neurophysiological signals of working memory in normal aging. Cognitive Brain Research, 11(3), 363–376. 10.1016/S0926-6410(01)00009-X

McKeown, D. J., Roberts, E., Finley, A. J., Kelley, N. J., Keage, H. A. D., Schinazi, V. R., Baumann, O., Moustafa, A. A., & Angus, D. J. (2025). Lower aperiodic EEG activity is associated with reduced verbal fluency performance across adulthood. Neurobiology of Aging, 151, 29–41. 10.1016/j.neurobiolaging.2025.03.013

Merkin, A., Sghirripa, S., Graetz, L., Smith, A. E., Hordacre, B., Harris, R., Pitcher, J., Semmler, J., Rogasch, N. C., & Goldsworthy, M. (2023). Do age-related differences in aperiodic neural activity explain differences in resting EEG alpha? Neurobiology of Aging, 121, 78–87. 10.1016/j.neurobiolaging.2022.09.003

Neubauer, A. C., & Fink, A. (2009). Intelligence and neural efficiency. Neuroscience & Biobehavioral Reviews, 33(7), 1004–1023. 10.1016/j.neubiorev.2009.04.001

Osaka, N., Logie, R. H., & D’Esposito, M. (2007). The Cognitive Neuroscience of Working Memory. Oxford University Press.

Podvalny, E., Noy, N., Harel, M., Bickel, S., Chechik, G., Schroeder, C. E., Mehta, A. D., Tsodyks, M., & Malach, R. (2015). A unifying principle underlying the extracellular field potential spectral responses in the human cortex. Journal of Neurophysiology, 114(1), 505–519. 10.1152/jn.00943.2014

Polich, J. (2007). Updating P300: An integrative theory of P3a and P3b. Clinical Neurophysiology, 118(10), 2128–2148. 10.1016/j.clinph.2007.04.019

Prince, J. B., Davis, H. L., Tan, J., Muller-Townsend, K., Markovic, S., Lewis, D. M. G., Hastie, B., Thompson, M. B., Drummond, P. D., Fujiyama, H., & Sohrabi, H. R. (2024). Cognitive and neuroscientific perspectives of healthy ageing. Neuroscience & Biobehavioral Reviews, 161, 105649. 10.1016/j.neubiorev.2024.105649

Proskovec, A. L., HeinrichsDGraham, E., & Wilson, T. W. (2016). Aging modulates the oscillatory dynamics underlying successful working memory encoding and maintenance. Human Brain Mapping, 37(6), 2348–2361. 10.1002/hbm.23178

Reuter-Lorenz, P. A., & Cappell, K. A. (2008). Neurocognitive Aging and the Compensation Hypothesis: Current Directions in Psychological Science. http://journals.sagepub.com/doi/10.1111/j.1467-8721.2008.00570.x

Salthouse, T. A. (1994). The aging of working memory. Neuropsychology, 8(4), 535–543. 10.1037/0894-4105.8.4.535

Sghirripa, S., Graetz, L., Merkin, A., Rogasch, N. C., Semmler, J. G., & Goldsworthy, M. R. (2021). Load-dependent modulation of alpha oscillations during working memory encoding and retention in young and older adults. Psychophysiology, 58(2), e13719. 10.1111/psyp.13719

Smith, A. E., Chau, A., Greaves, D., Keage, H. A. D., & Feuerriegel, D. (2023). Resting EEG power spectra across middle to late life: Associations with age, cognition, APOE-D4 carriage, and cardiometabolic burden. Neurobiology of Aging, 130, 93–102. 10.1016/j.neurobiolaging.2023.06.004

Snodgrass, J. G., & Corwin, J. (1988). Pragmatics of measuring recognition memory: Applications to dementia and amnesia. Journal of Experimental Psychology: General, 117(1), 34–50. 10.1037/0096-3445.117.1.34

Solé-Padullés, C., Bartrés-Faz, D., Junqué, C., Vendrell, P., Rami, L., Clemente, I. C., Bosch, B., Villar, A., Bargalló, N., Jurado, M. A., Barrios, M., & Molinuevo, J. L. (2009). Brain structure and function related to cognitive reserve variables in normal aging, mild cognitive impairment and Alzheimer’s disease. Neurobiology of Aging, 30(7), 1114–1124. 10.1016/j.neurobiolaging.2007.10.008

Speer, M. E., & Soldan, A. (2015). Cognitive reserve modulates ERPs associated with verbal working memory in healthy younger and older adults. Neurobiology of Aging, 36(3), 1424–1434. 10.1016/j.neurobiolaging.2014.12.025

Springer, S. D., Okelberry, H. J., Willett, M. P., Johnson, H. J., Meehan, C. E., Schantell, M., Embury, C. M., Rempe, M. P., & Wilson, T. W. (2023). Age-related alterations in the oscillatory dynamics serving verbal working memory processing. Aging (Albany NY), 15(24), 14574–14590. 10.18632/aging.205403

Stern, Y. (2009). Cognitive reserve. Neuropsychologia, 47(10), 2015–2028. 10.1016/j.neuropsychologia.2009.03.004

Stevens, W. D., Hasher, L., Chiew, K. S., & Grady, C. L. (2008). A Neural Mechanism Underlying Memory Failure in Older Adults. Journal of Neuroscience, 28(48), 12820–12824. 10.1523/JNEUROSCI.2622-08.2008

Thuwal, K., Banerjee, A., & Roy, D. (2021). Aperiodic and Periodic Components of Ongoing Oscillatory Brain Dynamics Link Distinct Functional Aspects of Cognition across Adult Lifespan. eNeuro, 8(5). 10.1523/ENEURO.0224-21.2021

Tran, T. T., Rolle, C. E., Gazzaley, A., & Voytek, B. (2020). Linked Sources of Neural Noise Contribute to Age-related Cognitive Decline. Journal of Cognitive Neuroscience, 1–110. 10.1162/jocn_a_01584

Vehtari, A., Gelman, A., Simpson, D., Carpenter, B., & Bürkner, P.-C. (2021). Rank-Normalization, Folding, and Localization: An Improved R^ for Assessing Convergence of MCMC (with Discussion). Bayesian Analysis, 16(2), 667–718. 10.1214/20-BA1221

Virtue-Griffiths, S., Fornito, A., Thompson, S., Biabani, M., Tiego, J., Thapa, T., & Rogasch, N. C. (2022). Task-related changes in aperiodic activity are related to visual working memory capacity independent of event-related potentials and alpha oscillations (p. 2022.01.18.476852). bioRxiv. 10.1101/2022.01.18.476852

Voytek, B., & Knight, R. T. (2015). Dynamic Network Communication as a Unifying Neural Basis for Cognition, Development, Aging, and Disease. Biological Psychiatry, 77(12), 1089–1097. 10.1016/j.biopsych.2015.04.016

Voytek, B., Kramer, M. A., Case, J., Lepage, K. Q., Tempesta, Z. R., Knight, R. T., & Gazzaley, A. (2015). Age-Related Changes in 1/f Neural Electrophysiological Noise. The Journal of Neuroscience, 35(38), 13257–13265. 10.1523/JNEUROSCI.2332-14.2015

Waschke, L., Donoghue, T., Fiedler, L., Smith, S., Garrett, D. D., Voytek, B., & Obleser, J. (2021). Modality-specific tracking of attention and sensory statistics in the human electrophysiological spectral exponent. eLife, 10, e70068. 10.7554/eLife.70068

Waschke, L., Wöstmann, M., & Obleser, J. (2017). States and traits of neural irregularity in the age-varying human brain. Scientific Reports, 7(1), 17381. 10.1038/s41598-017-17766-4

Wickham, Hadley. (2016). Ggplot2: Elegant graphics for data analysis (2nd ed.). Springer International Publishing.

Zarahn, E., Rakitin, B., Abela, D., Flynn, J., & Stern, Y. (2007). Age-related changes in brain activation during a delayed item recognition task. Neurobiology of Aging, 28(5), 784– 798. 10.1016/j.neurobiolaging.2006.03.002

Zhang, C., Stock, A.-K., Mückschel, M., Hommel, B., & Beste, C. (2023). Aperiodic neural activity reflects metacontrol. Cerebral Cortex, 33(12), 7941–7951. 10.1093/cercor/bhad089

